# Life-long hydroxyurea treatment decreases chronic pain in sickle cell disease mice

**DOI:** 10.1101/2025.08.30.673263

**Authors:** Nyabouk J. Gayluak, Sourav S. Roy, Ashley N. Plumb, Hemanth Mydugolam, Zulmary Manjarres, Lindsay E. Ramos Freitas, Amanda M. Brandow, Michael D. Burton, Katelyn E. Sadler

**Affiliations:** Department of Neuroscience, Center for Advanced Pain Studies, University of Texas at Dallas; Richardson, TX, USA; Department of Pediatrics, Division of Hematology/Oncology/Bone Marrow Transplantation, Medical College of Wisconsin; Milwaukee, WI, USA

## Abstract

Chronic sickle cell disease (SCD) pain mechanisms remain critically understudied, even though more than 50% of patients develop this symptom as their disease progresses. Despite high face validity, there are critical gaps in transgenic SCD mouse model characterization and implementation that must be addressed in order to increase the translational relevance of these animals. First, it is unclear when the chronic pain phenotype first develops in these mice. Second, there are no studies that have measured chronic pain in animals following standard-of-care drug regimens. Herein, we address both of these gaps by performing reflexive pain behavior tests in hydroxyurea-treated Townes HbSS and HbAA mice from postnatal day 10 to 6 months of age. Hydroxyurea (HU), a compound that increases circulating levels of fetal hemoglobin (HbF), is a life-long therapy prescribed to individuals with SCD beginning as early as age 9 months. Here, we demonstrate that chronic mechanical hypersensitivity develops in Townes HbSS mice between P21-P28, a time frame that follows the HbF-to-HbS switch. When initiated at birth, HU treatment limits the extent of chronic mechanical pain development in HbSS mice. HU analgesic effects can be attributed to decreased innate immune tone in the periphery; life-long HU treatment decreases circulating monocyte counts in HbSS mice and reverses sensitization of TRPA1, a lipopolysaccharide receptor, in HbSS nociceptors. In conclusion, these studies provide additional support for early implementation of HU in SCD disease management, and furthermore, identify the LPS-TRPA1 signaling axis as a novel analgesic target for chronic SCD pain.

## Introduction

Pain is the leading reason individuals with SCD seek medical attention^1^. In addition to suffering from acute sickle cell “pain crises” – experiences that are widely attributed to vasoocclusion – over 50% of individuals with SCD also experience chronic pain that has no clear etiology^2^. Chronic SCD pain is frequently managed with opioid drugs. For individuals with SCD, these therapies have unwanted side effects, they provide incomplete analgesia, and they are often challenging to procure^3^. Thus, additional treatments for chronic SCD pain are desperately needed. To this end, the landmark approval of the first two SCD gene therapies – exa-cel and lovo-cel – were met with great enthusiasm; surely improvement in disease pathophysiology would also decrease chronic pain in patients. However, notably omitted from the associated clinical trials were chronic pain measures^4,5^. While waiting to make conclusions about the efficacy of these therapies, inferences can be drawn from previous reports of individuals who received hematopoietic stem cell transplant (HSCT). Numerous studies suggest that following HSCT, a subset of individuals who previously had SCD will continue to suffer from chronic pain^6–8^. Collectively, these data suggest that early disease management may be critical for minimizing pathophysiology and preventing the centralization of pain in individuals with SCD. Thus, to develop the best analgesics for this patient population, we need to first understand when chronic pain develops in SCD and secondarily determine if life-long disease modifying therapies alters chronic pain trajectories.

Hydroxyurea (HU) is a neoplastic agent that was adopted as a SCD therapy after it was determined to induce the expression of fetal hemoglobin (HbF)^9,10^. Unlike the mutated version of adult hemoglobin (HbS), HbF does not polymerize. By increasing production of HbF in individuals with SCD, HU decreases the likelihood of red blood cell (RBC) sickling. Initial clinical trials and years of subsequent study show that acute (< 6 months) HU treatment improves hematological parameters^11^, decreases the rate of painful crises^11^, decreases daily pain intensity^12^, and decreases analgesic use^12^ in adults with SCD. Extended HU use also decreases the frequency of acute SCD pain episodes^13^, and both heat and mechanical pain thresholds in adult patients^14^. Collectively, these studies provide strong support for HU improving acute SCD pain metrics. Less clear however, is what effect HU has on chronic SCD pain metrics, particularly when the treatment regimen is initiated in infancy. Thus, in these studies, we employed the Townes model to determine when chronic pain first develops in SCD mice, and to subsequently assess if life-long hydroxyurea treatment decreases chronic pain measures.

## Materials and Methods

### Animals

Male and female Townes HbAA (control) and HbSS (SCD) mice were used for all experiments^15^. Townes mice express human α, human γ, and human β (A or S) globin genes instead of endogenous mouse globins. To generate HbSS mice, HbAS dams were bred with HbSS studs; HbAS offspring were culled. To generate AA mice, HbAA dams were bred with HbAA studs. Animals were maintained on a 12-hour light/dark cycle and experiments were performed during the light cycle. All animals had access to *ad libitum* food and water. All protocols were in accordance with National Institutes of Health guidelines and were approved by the Institutional Animal Care and Use Committee at the University of Texas at Dallas (protocol 2022-0088).

### Western blot

To determine when Townes mice stop expressing γ-globin protein, western blots were performed as previously described^16^. In short, blood samples were collected from mice at various ages, then used for western blot comparisons of γ-globin protein levels. γ-globin was normalized to total globin in each sample.

### *In vitro* sickling assay

Blood samples were collected from mice of various ages or following 6 months of hydroxyurea treatment then exposed to 0.2% sodium metabisulfite *in vitro* to induce sickling as previously described^17^. Sickled RBCs (as a % of total blood cells) were counted by an experimenter blinded to treatment and genotype.

### Hydroxyurea (HU) treatment

All mice were randomized to treatment group. HU treatment was administered at 50 mg/kg/day via drinking water to Townes dams starting on the day pups were born (P0)^18^. Pups received HU treatment indirectly through milk. Following weaning at P21, mice continued to receive the clinically appropriate dose of 25 mg/kg/day HU via drinking water until the time of euthanasia^19^. To ensure effects of HU could not be attributed to growth delays, all animals were weighed monthly. No significant differences in body weight were observed between treatment groups (data not shown). Dams were euthanized following weaning to prevent issues with future litters given that HU can be teratogenic and embryotoxic^20^.

### Behavior tests

Animals younger than P21 were habituated and tested for a maximum of 40 min to limit stress and time away from dams. Mice aged P21 and older were habituated for 2 hr and tested thereafter. The experimenter was present in the testing room for 30 min before starting all experiments to allow for olfactory signal habituation^21^. The behavior room was maintained on average at 22.0 °C, equipped with a white noise machine and overhead white lights remained on throughout the experiment. The experimenter was blinded to the treatment and genotype in all experiments.

#### Up-down von Frey mechanical sensitivity testing

To assess age-dependent punctate mechanical allodynia, von Frey testing was performed as previously described on the dorsal hindpaw for animals <P21^22^ and the plantar surface of the hindpaw for animals >P21^23^. 50% withdrawal thresholds were calculated for each paw as previously described^24^ then averaged between paws.

#### Dynamic brush mechanical sensitivity testing

To assess age-dependent dynamic mechanical allodynia, a 10/0 short liner paintbrush was placed directly above (< P21) or underneath (>P21) each hindpaw then gently swept from heel to toe. Animal reactions were qualified as null (no response), normal (paw was lifted and immediately returned to floor), or nocifensive (slamming, shaking, guarding licking or any additional attending to paw). Brush stimulation was repeated five times on each hindpaw. Response types from both paws were summed for each animal.

#### Needle mechanical sensitivity testing

To assess age-dependent punctate mechanical hyperalgesia, a 27-gauge needle was used to poke the center of each hindpaw (dorsal surface for < P21, plantar surface for >P21). Each paw was stimulated five times; responses were characterized and summed as in the brush test.

#### Planar dry ice-cold sensitivity testing

The plantar dry ice test was performed in animals of all ages as previously described. Withdrawal latencies were recorded five times for each paw then averaged for a given animal^25^.

#### TRPA1 antagonist experiments

HbSS mice were randomly assigned to receive either vehicle (saline) or 100 mg/kg HC-030031 i.p.^26^. von Frey thresholds were assessed 1.5 hr after treatment.

### Blood parameters and metabolite analysis

At 6 months of age, blood was collected from all animals via cardiac puncture. Hematological parameters were assessed on whole blood using the VetScan HM5 Hematology System (Zoetis). Plasma metabolites were measured using VetScan Preventative Care Profile Plus rotors (Zoetis).

### *In vitro* calcium imaging

Bilateral dorsal root ganglia (DRG; L1–L6) were isolated and cultured from 6 month old HbAA and HbSS mice according to established protocols from our laboratory^27,28^. 24hr after isolation, neurons were loaded with the dual-wavelength ratiometric calcium indicator Fura-2AM then imaged while being exposed to lipopolysaccharide (LPS, 10, 50, or 100 µg/mL; 3 min exposure); neuron health was confirmed by examining responses to 50 mM KCl. Fura-2 emissions were analyzed using a custom-designed script. In TRPA1 antagonist experiments, DRG neurons were incubated in 10 µM HC-030031 (dissolved in 0.1% DMSO) for 15 minutes at room temp prior to LPS exposure.

### Statistical analysis

Data were analyzed using GraphPad Prism 10. Results were considered statistically significant when *P* < 0.05. Data are presented as group means ± SEM.

## Results

### Townes HbSS mice undergo fetal-to-adult hemoglobin switch during late embryonic development

Before assessing the effects of hydroxyurea (HU) on persistent sickle cell disease (SCD) pain, we first needed to identify the timepoint when mechanical and thermal hypersensitivity develops in transgenic SCD mice. Individuals with SCD do not exhibit chronic pain at birth. Rather, chronic pain develops over time following the fetal-to-adult hemoglobin switch which occurs in the first year of life. During this time, decreases in fetal hemoglobin (HbF) expression are paralleled by increased expression of either adult hemoglobin (HbA) in healthy individuals or the mutated hemoglobin variant (HbS) in those with SCD (**Figure 1A**). Since HbF does not polymerize, we hypothesized that, like individuals with SCD, Townes HbSS mice would not develop persistent pain before the HbF-to-HbS switch. To pinpoint the timing of the HbF-to-HbS transition in Townes HbSS mice, western blot analysis was used to measure relative γ-globin (subunit of HbF) levels in blood collected at various ages (**Figure 1B**). In blood samples collected on postnatal (P) days 0 and 2, the γ-to-total globin ratio was significantly higher than in samples collected from adult mice (**Figure 1C**). By P7 however, significant decreases in circulating HbF were observed. Given that the average lifespan of an erythrocyte in SCD mice is approximately 5 days^29^, this suggests that the fetal-to-adult hemoglobin switch occurs on or before P2, leading to markedly reduced HbF levels in circulation by P7. To confirm this timing, blood samples from Townes HbSS mice were mixed with sodium metabisulfite to induce sickling *in vitro*. Sickled red blood cells (RBCs) were observed in blood as early as P0 (**Figure 1D**) and sickling frequency increased with age (**Figure 1E**). Collectively, these data suggest that the epigenetic switch from HbF to HbS occurs during the final stages of embryonic development.

**Figure 1.**
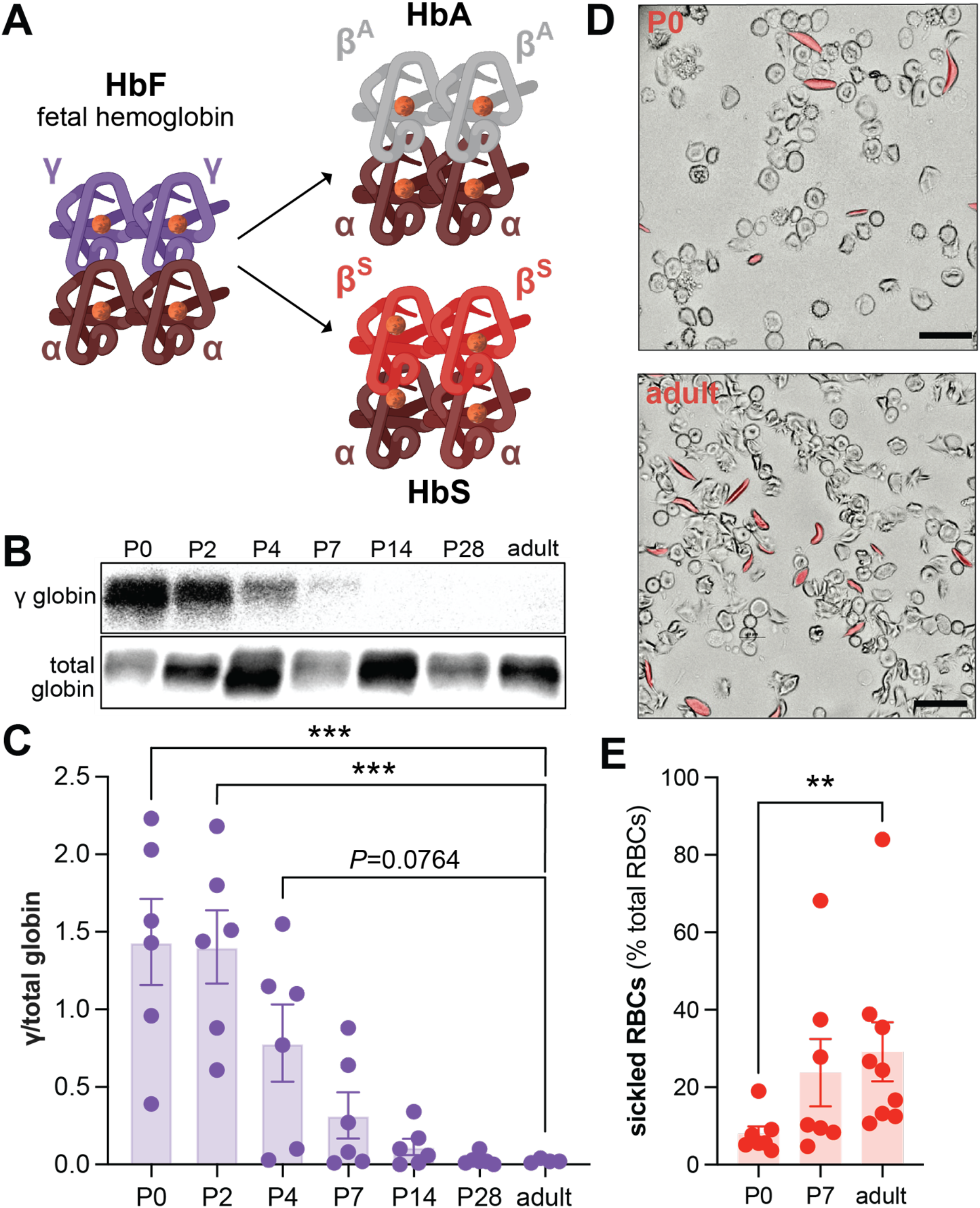
Time course of fetal-to-adult hemoglobin switch in Townes SCD mice. (**A**) Graphic representation of fetal (HbF) to HbA (adult) or HbS (sickle) hemoglobin switch. (**B**) Representative western blot depicting γ-globin and total globin blood proteins at various timepoints in Townes SCD mice. Adult samples were from 7-month-old mice. (**C**) The relative expression of γ-globin to total globin in circulation with increasing age in mice (1-way ANOVA: *P*<0.001, Bonferroni’s multiple comparison: \*\*\**P*<0.001). (**D**) Representative images of *in vitro* sickling assay performed on blood collected from P0 and 6-month-old adult mice. Bars represent 20µm. (**E**) Quantification of sickled red blood cells (RBCs) following *in vitro* sickling assay (Kruskal Wallis test main effect *P*=0.0110, Bonferroni’s multiple comparison **P<0.01).

### Persistent mechanical hypersensitivity develops in Townes HbSS mice between postnatal days 21 and 28

Having established that RBCs in Townes HbSS blood can sickle by P0 and that circulating HbF levels are significantly decreased by P7, we hypothesized that mechanical and thermal hypersensitivity would be observed after P7. To test this, sensitivity to evoked mechanical and thermal stimuli was measured in Townes HbSS and HbAA mice starting at P10. Punctate mechanical allodynia (*i.e*., hypersensitivity to static light touch) was first measured by applying calibrated von Frey filaments to the dorsal hairy surface (P10, P14) or ventral glabrous surface (P21, P28) of mouse hindpaws. By P28, Townes HbSS mice exhibited significantly lower withdrawal thresholds than Townes HbAA mice, indicating that mechanical hypersensitivity was established by this age (**Figure 2A**). Dynamic mechanical allodynia (*i.e*., hypersensitivity to kinetic light touch) was assessed by dragging a fine-tipped paintbrush across mouse hindpaws. Brush stimulation of the dorsal hindpaw elicited comparable responses in HbSS and HbAA mice at P10 and P14 (**Figure 2B**). By P21, HbSS mice exhibited more nocifensive responses than HbAA control mice indicating that dynamic mechanical allodynia developed by this age. Finally, punctate mechanical hyperalgesia (*i.e*., exaggerated sensitivity to static painful touch) was assessed by stimulating hindpaws with a needle. Needle stimulation of the dorsal hindpaw elicited similar responses in HbSS and HbAA mice at P10 and P14 (**Figure 2C**). However, by P21, HbSS mice exhibited hypersensitivity to needle stimulation as evidenced by both a decrease in null responses and an increase in nocifensive responses. Thus, mechanical behavior tests revealed that HbSS mice develop hypersensitivity to needle and brush stimuli by P21 and static mechanical allodynia by P28.

**Figure 2.**
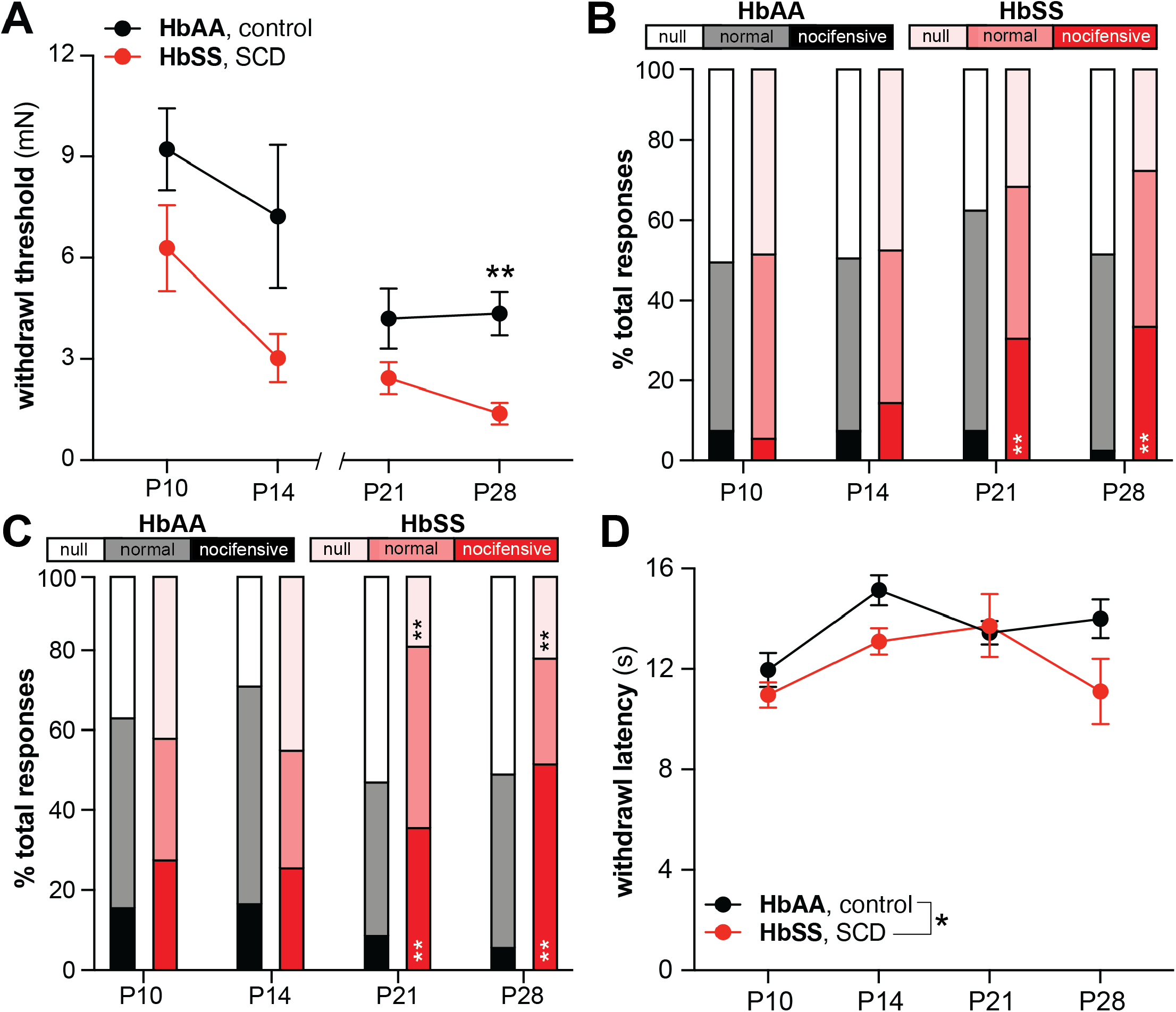
Time course of persistent pain development in Townes SCD mice. (**A**) Hindpaw mechanical withdrawal thresholds of Townes HbAA (control) and HbSS (SCD) mice at various ages (P10 and P14: *N*=6-16; 2-way ANOVA, main effect of genotype: *P*=0.0123; P21 and P28: *N*=10-14; 2-way ANOVA, main effect of genotype: *P*=0.0081, Bonferroni’s multiple comparison: \*\**P*=0.0025). (**B**) Responses to hindpaw dynamic brush stimulation over first month of life (*N*=6-16; Chi-square overall analysis *P*<0.05; corrected Fisher’s exact multiple comparisons AA vs. SS ** *P*<0.01). (**C**) Responses to hindpaw needle stimulation over first month of life (*N*=6-16; Chi-square overall analysis *P*<0.05; corrected Fisher’s exact multiple comparisons AA vs. SS ** *P*<0.01). (**D**) Hindpaw cold withdrawal latency of Townes HbAA and HbSS mice over first month of life (*N*=4-10; 2-way ANOVA, main effect of genotype: \**P*=0.0031, of age: *P*=0.0217).

Given that adult humans and mice with SCD also exhibit robust cold hypersensitivity^27,30,31^, we next used the plantar dry ice test to determine if this phenotype also exists in younger (P0-P28) HbSS mice. Although there was a main effect of genotype in our statistical analysis, post-hoc tests did not yield significant differences between HbSS and HbAA mice at individual time points tested (**Figure 2D**). Therefore, these findings suggest that robust cold hypersensitivity develops later than mechanical hypersensitivity in Townes HbSS mice.

### Life-long hydroxyurea treatment reduces mechanical hypersensitivity in Townes SCD mice

After establishing that mechanical hypersensitivity develops in HbSS mice between P21 and P28 (roughly equivalent to late juvenile/early peripubertal stages in human)^32^, we next examined whether this phenotype could be reversed by life-long hydroxyurea (HU) treatment. To this end, new cohorts of HbAA and HbSS mice were enrolled in either HU or vehicle treatment on P0. Subsequent changes in evoked mechanical and thermal sensitivity were evaluated over the first 6 months of life. Similar to our original findings (**Figure 2A**), vehicle-treated HbSS mice exhibited static touch hypersensitivity by P28 as compared to vehicle-treated HbAA mice (**Figure 3A**). Notably, this mechanical hypersensitivity phenotype was reduced in HU-treated HbSS as early as P28. In contrast, HU-treated HbAA mice developed mechanical hypersensitivity over time. This paradoxical effect may reflect the chemotherapeutic properties of HU. HU use has been linked to peripheral neuropathy^33^, a condition which could account for the increased mechanical sensitivity in control animals. In summary, life-long HU treatment significantly reduced the punctate mechanical allodynia phenotype exhibited by Townes HbSS mice and induced a state of chronic touch hypersensitivity in HbAA control mice (**Figure 3B**).

**Figure 3.**
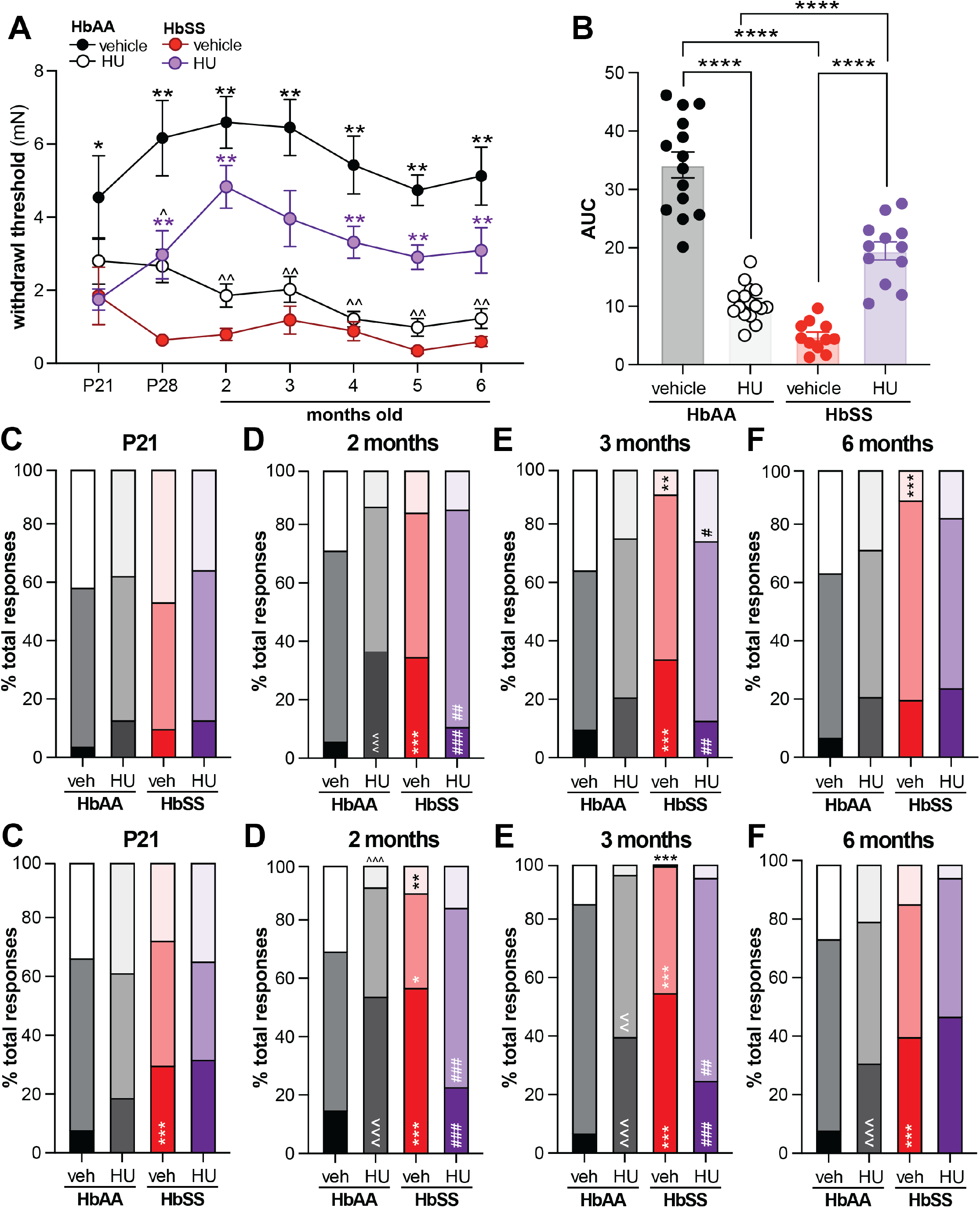
Long-term hydroxyurea treatment reduces chronic touch hypersensitivity in Townes SCD mice. (**A**) Hindpaw mechanical withdrawal thresholds of Townes HbAA (control) and HbSS (SCD) mice at various time points during hydroxyurea (HU) or vehicle treatment (*N*=9-12; mixed-effects analysis, main effect of genotype: *P*<0.0001, of drug: *P*=0.0079, genotype x drug interaction: *P*<0.0001, genotype x time x drug interaction *P*=0.0049; Bonferroni’s multiple comparisons: AA-vehicle vs. SS-vehicle \**P*<0.05, \*\**P*<0.01, AA-vehicle vs. AA-HU ^*P*<0.05, ^^*P*<0.01, SS-vehicle vs. SS-HU purple \**P*<0.05, \*\**P*<0.01). (**B**) Area under the curve (AUC) analysis of hindpaw von Frey thresholds over 6-month vehicle or HU-treatment (2-way ANOVA, main effect of treatment *P*=0.0042, genotype *P*<0.0001, genotype x treatment interaction *P*<0.000; Bonferroni’s multiple comparisons: \*\*\*\**P*<0.0001). Hindpaw responses to dynamic brush stimulus at (**C**) P21, (**D**) 2 months, (**E**) 3 months, and (**F**) 6 months of age (*N*=9-12; corrected Fisher’s exact multiple comparisons AA-vehicle vs. SS-vehicle ** *P*<0.01, \*\*\**P*<0.001, AA-vehicle vs. AA-HU ^^^*P*<0.001, SS-vehicle vs. SS-HU #*P*<0.05, ##*P*<0.01, ###*P*<0.01). Hindpaw responses to needle stimulus at (**G**) P21, (**H**) 2 months, (**I**) 3 months, and (**J**) 6 months of age (*N*=9-12; corrected Fisher’s exact multiple comparisons AA-vehicle vs. SS-vehicle ** *P*<0.01, \*\*\**P*<0.001, AA-vehicle vs. AA-HU ^^*P*<0.01, ^^^*P*<0.001, SS-vehicle vs. SS-HU ##*P*<0.01, ###*P*<0.01).

HU effects on dynamic mechanical allodynia were next assessed using the brush assay. Like previous results, vehicle-treated HbSS mice exhibited either more nocifensive responses or fewer null responses than vehicle-treated HbAA mice beginning at 2 months of age (**Figures 3C-3F**). HU treatment reversed this allodynia in HbSS mice, specifically at 2 and 3 months of age. By 6 months, HU-treated HbSS mouse responses to brush were identical to responses exhibited by vehicle-treated HbSS mice (**Figure 3F**). Similar time-dependent HU effects were also observed in the needle test. Vehicle-treated HbSS mice exhibited mechanical hyperalgesia beginning as early as P21 (**Figure 3G**), but HU-treated HbSS mice only exhibited a reversal of this phenotype at 2 and 3 months of age (**Figure 3G-3J**). In summary, life-long HU treatment alleviates chronic touch pain throughout the lifespan and reverses kinetic light touch pain and mechanical hyperalgesia in an age-specific manner.

### Life-long hydroxyurea treatment improves hematological parameters and reduces erythrocyte sickling in Townes SCD mice

Having established that HU treatment reduces chronic SCD mechanical hypersensitivity, we next sought to investigate the basis of this analgesia. HU is a primary SCD modifying therapy due to its ability to increase circulating HbF and reduce erythrocyte sickling, the products of which drive chronic SCD pain^34^. To determine if HU has similar effects in Townes SCD mice, hematological parameters were measured in blood collected from HbSS and HbAA mice following 6 months of HU or vehicle treatment. As expected, RBC count and total hemoglobin levels were lower in vehicle-treated HbSS mice compared to vehicle-treated controls. HU treatment did not reverse these anemia indicators in HbSS mice (**Figure 4A, 4B**). HU treatment did, however, reduce the elevated mean corpuscular volume (MCV) observed in HbSS animals (**Figure 4C**). This result supports the idea that HU treatment leads to a higher proportion of healthy, normally sized cells in circulation. Finally, and perhaps most notably, *in vitro* exposure to sodium metabisulfite revealed fewer sickled erythrocytes in blood collected from HU-treated HbSS mice as compared with vehicle-treated HbSS mice (**Figure 4D, 4E**). Paralleling this finding, significantly less bilirubin, a byproduct of hemolysis, was observed in blood from HU-treated HbSS mice as compared to vehicle-treated HbSS blood (**Figure 4F**). Collectively, these results support the observation that chronic HU treatment reduces pathophysiology in Townes HbSS mice.

**Figure 4.**
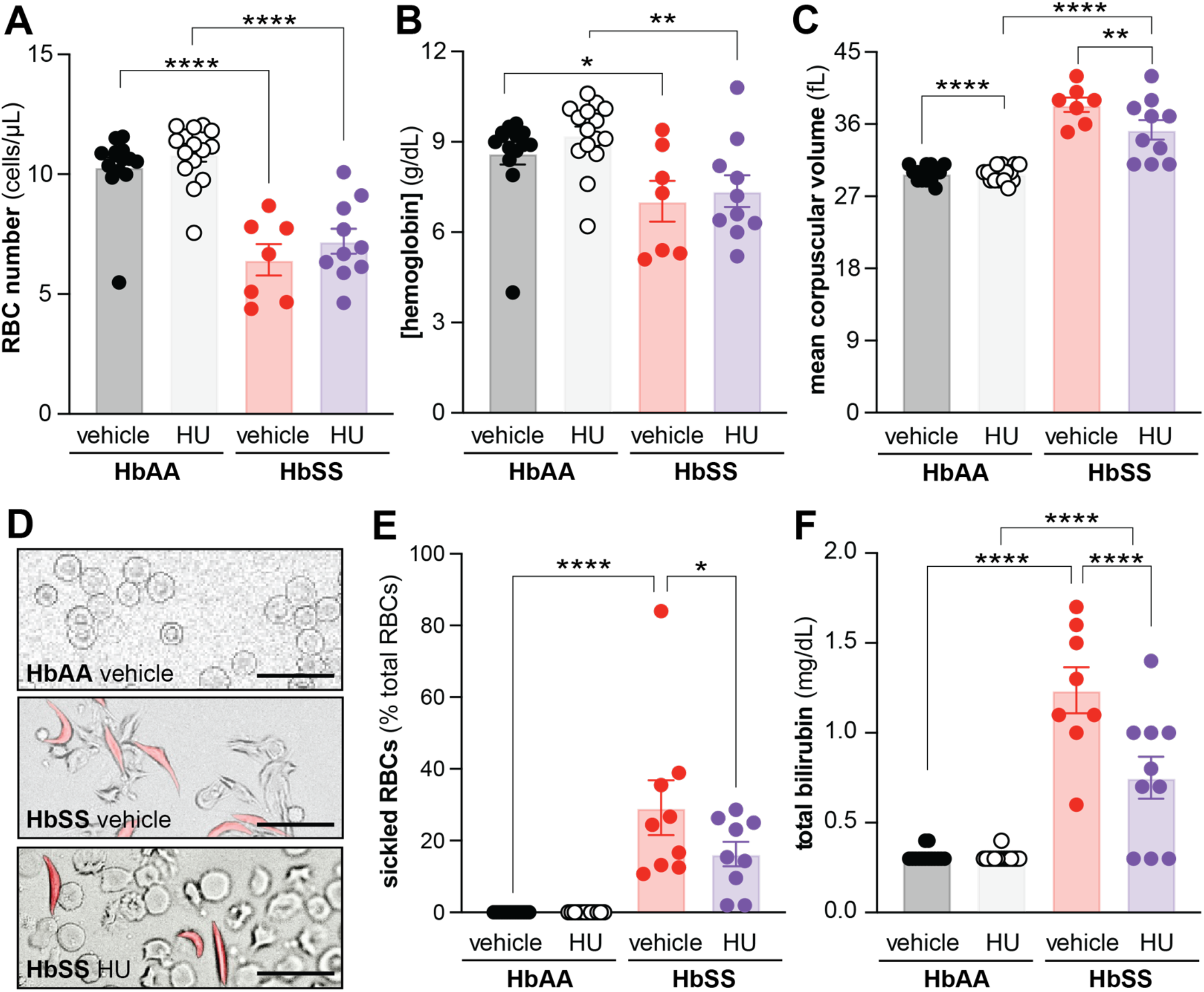
Life-long hydroxyurea treatment alters hematological parameters in Townes SCD mouse blood. Blood was collected from Townes HbAA and HbSS mice following 6 months of vehicle or hydroxyurea treatment. The following hematological parameters were recorded: (**A**) red blood cell count (2-way ANOVA main effect of genotype *P*<0.0001) (**B**) hemoglobin concentration (2-way ANOVA main effect of genotype *P*=0.0004) (**C**) mean corpuscular volume (2-way ANOVA main effect of genotype *P*<0.0001, treatment *P*=0.0202, genotype x treatment interaction *P*=0.0248). **(D)** Representative images of sodium metabosulfite exposed blood from vehicle-treated HbAA mice, vehicle- and HU-treated HbSS mice. Bar represents 20µm. (**E**) Percentage of all red blood cell that sickle in *in vitro* hypoxia assay. (2-way ANOVA main effect of genotype: *P*<0.0001). (**F**) Total bilirubin levels in blood of HbAA or HbSS mice following 6 months of vehicle or hydroxyurea treatment (2-way ANOVA main effect of treatment *P*=0.0007, genotype *P*<0.0001, treatment x genotype interaction *P*<0.0001). For all panels, Bonferroni’s multiple comparisons \**P*<0.05, \*\**P*<0.01, \*\*\**P*<0.001.

### Hydroxyurea analgesia results from decreased TRPA1-dependent neuronal sensitization to lipopolysaccharide

Chronic inflammation is another hallmark of SCD^34,35^. To determine if HU-associated analgesia in HbSS mice is associated with a reduction in this inflammation, multiple tissues and cell types were examined. Erythrocyte congestion, bone-marrow independent hematopoiesis, and lymphocyte accumulation collectively lead to splenomegaly in Townes HbSS mice^36^. This phenotype was observed in vehicle-treated HbSS animals and unchanged in the majority of HU-treated HbSS mice (**Figures 5A, 5B**). However, sub-sets of HU-treated HbSS mice exhibited further exacerbation or reversal of this phenotype suggesting that chronic HU treatment can have divergent effects, even when administered to a homogeneous population. Although HU treatment did not reduce increased white blood cell (**Figure 5C**) or lymphocyte (**Figure 5D**) counts in HbSS mice, it led to significant decreases in circulating monocyte number (**Figure 5E**).

**Figure 5.**
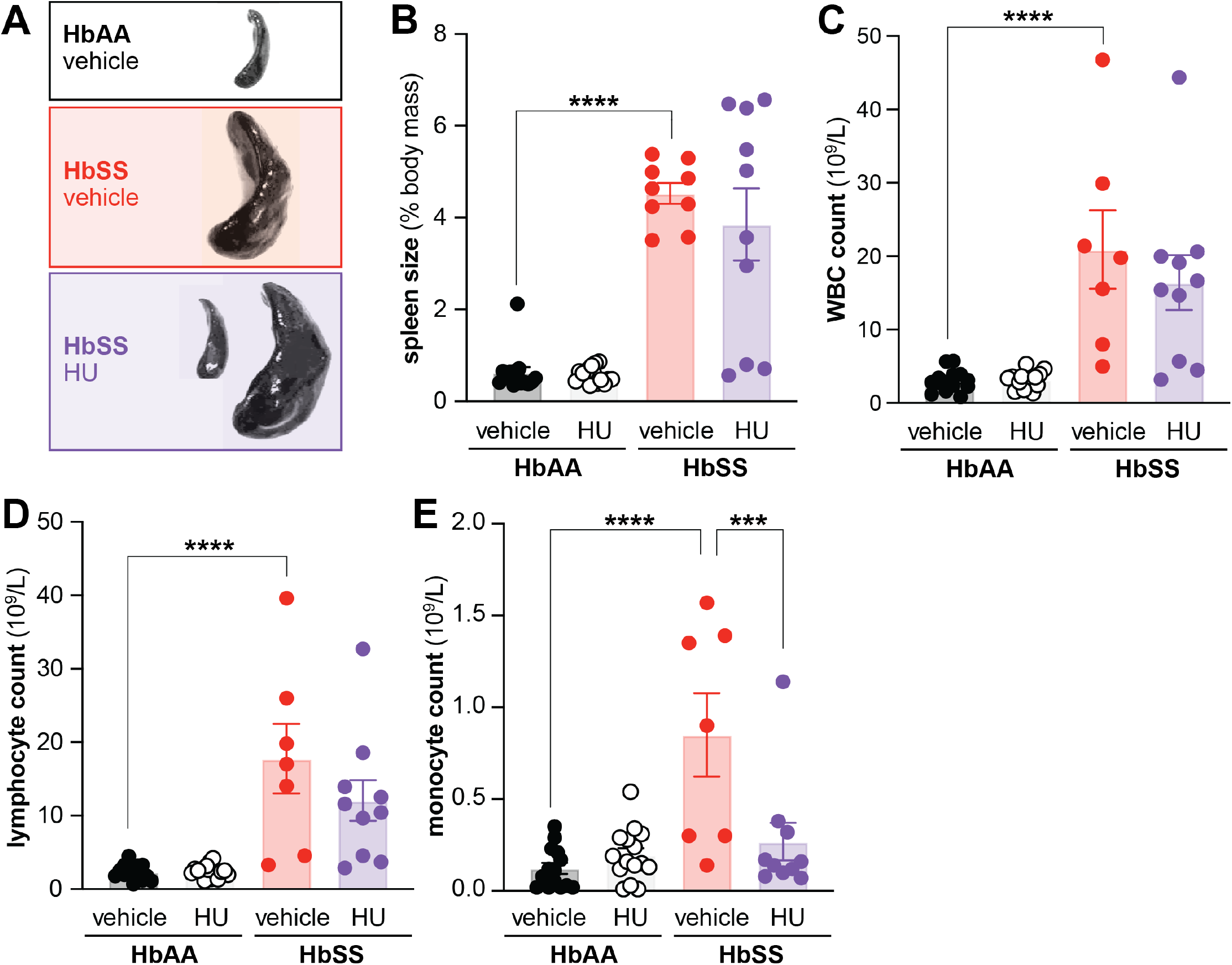
Life-long hydroxyurea treatment reduces innate immune response in Townes SCD mice. (**A**) Representative spleens isolated from vehicle treated HbAA and HbSS mice and hydroxyurea (HU)-treated HbSS mice. The following immune-related parameters were recorded in HbAA and HbSS mice following 6 months of vehicle or HU treatment: (**B**) relative spleen mass (2-way ANOVA main effect of genotype *P*<0.0001), (**C**) circulating white blood cell count (2-way ANOVA main effect of genotype *P*<0.0001), (**D**) circulating lymphocyte count (2-way ANOVA main effect of genotype *P*<0.0001) and (**E**) circulating monocyte count (2-way ANOVA main effect of genotype *P*<0.0001). For all panels, Bonferroni’s multiple comparisons \*\*\**P*<0.001, \*\*\**P*<0.0001.

Given the HU-associated reduction in innate immune system measures, HU-analgesia may ultimately result from decreased pathogen-associated molecular pattern (PAMP) signaling at the level of the peripheral nervous system. To directly assess this hypothesis, dorsal root ganglia (DRG) sensory neurons from vehicle-treated HbAA and HbSS mice were exposed to lipopolysaccharide (LPS), a component of Gram-negative bacteria outer membranes and a classic PAMP. Intracellular calcium flux was measured during extracellular LPS application to assess neuronal sensitivity to the agonist (**Figure 6A**). Although there was no genotype difference in the percentage of neurons that responded to application of 10 or 50 µg/mL LPS, significantly more HbSS DRG neurons exhibited calcium flux *(i.e*., indirect activity marker) in response to 100 µg/mL LPS than HbAA neurons (**Figure 6B**). This effect was observed in neurons >27 µm in diameter (**Figure 6C**) and in neurons < 27 µm in diameter (*i.e*., putative nociceptors; **Figure 6D**). Given that small diameter neurons are more likely to transmit nociceptive signals to the central nervous system, all subsequent analyses were only performed on these cells. Despite observed differences in proportion of activated neurons, HbAA and HbSS DRG neurons did not differ in terms of peak LPS-induced calcium response (**Figure 6E**), latency to peak response (**Figure 6F**), or extent of LPS response as measured by area under the curve analysis (**Figure 6G**). To determine if HU reverses LPS sensitization, calcium imaging was next performed on DRG isolated from HU-treated HbSS mice. Indeed, fewer neurons from HU-treated HbSS mice exhibited responses to LPS as compared to vehicle-treated HbSS mice (**Figure 6H**). LPS activity can be mediated by multiple receptors in neurons including toll-like receptor 4 (TLR4^37,38^) and the ionotropic receptor TRPA1^39^. Since TRPA1 is preferentially expressed in small diameter DRG neurons^40^, we next applied the TRPA1 antagonist HC-030031 to DRG neurons and measured subsequent responses to LPS application. HC-030031 reduced LPS responsivity of DRG neurons from HbAA, vehicle-treated HbSS, and HU-treated HbSS mice to such an extent that only ∼10% of neurons from each group remained responsive (**Figure 6I**). Given the statistical equivalence of remaining responders, we can thus assume that the exacerbated LPS responses in vehicle-treated HbSS mice were due to TRPA1 sensitization. Finally, to determine if TRPA1 activity contributes to chronic SCD pain i*n vivo*, mechanical sensitivity was assessed in vehicle-and HU-treated HbSS mice following HC-030031 administration. HC-030031 effectively alleviated chronic mechanical allodynia in vehicle-treated HbSS mice (**Figure 6J**) but had no effect in HU-treated HbSS animals (**Figure 6K**). In conclusion, HU analgesia results from blunted TRPA1 signaling in SCD primary sensory neurons.

**Figure 6.**
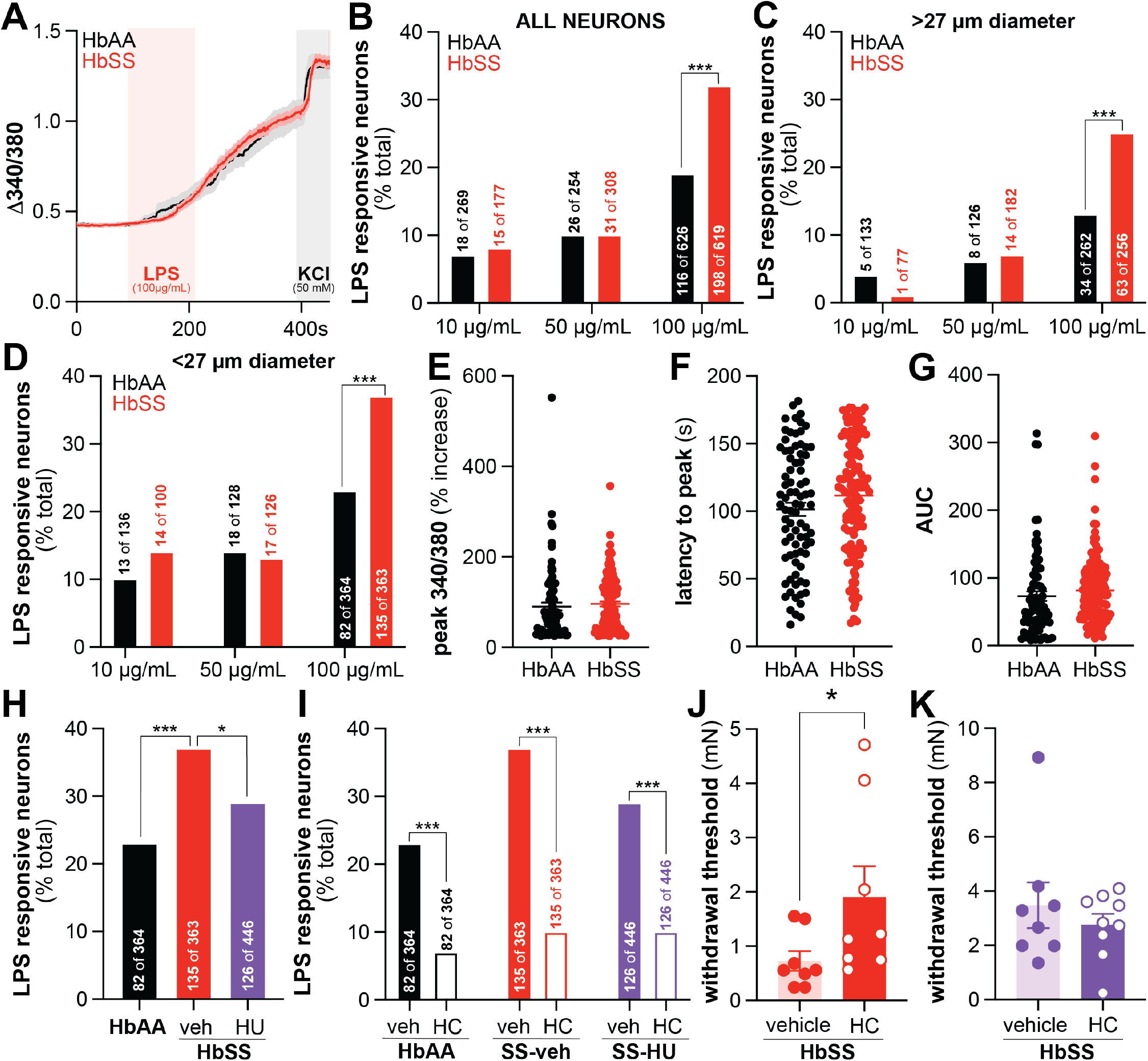
Life-long hydroxyurea treatment decreases SCD-associated LPS sensitization in dorsal root ganglia (DRG) sensory neurons. (**A**) Averaged intracellular calcium flux in DRG neurons isolated from HbAA and HbSS mice in response to extracellular application of LPS. Percentage of (**B**) all, (**C**) medium and large diameter, and (**D**) small diameter DRG neurons from HbAA and HbSS mice that exhibit increases in intracellular calcium upon extracellular exposure to increasing concentrations of LPS (corrected Fisher’s exact multiple comparisons \*\*\**P*<0.0001). Comparison of (**E**) peak 340/380 ratio, (**F**) latency to reach peak 340/380, and (**G**) 340/380 area under the curve from HbAA and HbSS small diameter DRG neurons during extracellular application of 100 µg/mL LPS. Percentage of small diameter DRG neurons that respond to extracellular application of 100 µg/mL LPS from (**H**) vehicle or HU-treated mice or (**I**) during co-incubation with the TRPA1 antagonist HC-030031 (10µM). Hindpaw withdrawal thresholds of (**J**) vehicle-treated and (**K**) HU-treated HbSS mice following i.p. injection of HC-030031 (100mg/kg).

## Discussion

Herein, we determined that life-long HU use decreases chronic mechanical hypersensitivity in SCD mice, in part, by preventing LPS sensitization of peripheral nociceptors. Decreased LPS neuronal responsivity in HU-treated SCD mice was paralleled by fewer circulating monocytes in the blood. Collectively, these data suggest that chronic HU-treatment limits the extent of innate immune surveillance that needs to occur in SCD mice. Heightened innate immune surveillance is correlated with chronic pain in individuals with SCD^41^. This phenotype may result from the increased bacterial infection susceptibility exhibited by both individuals and mouse models with SCD, or more likely, from the elevated levels of LPS in serum collected from both patients and SCD mice at steady state ^42,43^. Elevated serum LPS is attributed to increased gut permeability, a phenotype that is also reported in patients and mouse models with SCD ^42–45^. Although not directly assessed in this study, it is likely that HU-treated mice would exhibit decreased LPS levels in circulation, perhaps as an indirect result of reduced ischemic damage to the intestinal epithelium. In addition to reducing luminal accessibility, HU effects on circulating LPS could also result from drug-induced changes in the gut microbiota. Work from our lab and others has clearly demonstrated that gut dysbiosis drives chronic pain in SCD^46,47^. Early studies indicate that HU alters gut microbial composition in individuals with SCD^48^. Thus, in summary, HU analgesia may result from beneficial effects that aggregate at the level of the gastrointestinal tract; HU may both limit the extent of SCD-associated gut dysbiosis and prevent dysbiosis-associated metabolites from leaking out of the gut.

Many additional targets for chronic SCD pain are implicated by the results of these studies. First, it is obvious that an additional LPS receptor must be expressed by DRG neurons. The likely culprit for this activity is TLR4, a receptor that has previously been connected to chronic SCD pain due to its ability to directly bind heme^49^. In comparing LPS receptors, therapeutic targeting of TLR4 may prove more challenging than TRPA1 given its critical and widespread expression on myeloid-derived immune cells. Reducing TLR4 activity of the innate immune system could lead to further infection susceptibility in a clinical population already burdened by this difficulty. Second, it is likely that previously unstudied monocyte-secreted chemokines and cytokines also drive chronic SCD pain. For example, although our previous publication outlines a role for CCL2/CCR2 signaling in chronic SCD pain^27^, no preclinical studies have investigated the extent to which monocyte-derived TNF-α or IL-1? drive chronic SCD pain despite high levels of these compounds in steady state SCD circulation^50,51^ and HU-associated decreases in these compounts^52,53^. Indeed, administration of a TNF inhibitor for plaque psoriasis also resolved SCD pain in a single case report^54^ and, in a similar fashion, TNF blockade reduced pain in individuals suffering from both SCD and rheumatoid arthritis^12,14^. Thus, TNF is a high priority target, the analgesic efficacy of which should be systematically assessed in future preclinical studies.

Another notable finding of these studies is the time course of chronic pain development in transgenic Townes SCD mice. HbSS mice develop robust mechanical hypersensitivity between P21-P28, a period that roughly correlates with ages 8-12 in humans^32^. Although chronic pain is reported by subsets of individuals with SCD at these same ages^55^, retrospective studies have failed to identify molecular biomarkers that accurately predict which children will develop this symptom early – or ever – in their disease trajectory. Thus, we are now able to leverage the mouse model and the relatively short time scale over which pain develops in these animals to address this question systematically. The timing of mechanical hypersensitivity development in SCD mice is notable as it coincides with the period during which nociceptors undergo growth factor-based sub-class differentiation^56^, and nociceptor central terminals are pruned into relatively stable spinal cord circuits^57^. Despite exhibiting spinal reflexes in response to noxious stimulus application as early as P1^58^, these postnatal developments are required for directed and appropriate rodent behavioral responses to noxious stimulus application. Thus, it is possible that HbSS mice might exhibit exacerbated pain-like behaviors even earlier if only they were biologically equipped to do so. Our observation of RBC sickling – and theoretically, release of noxious contents from said cells – as early as P0 provide even further support for this idea. In addition to driving acute changes in sensitivity, this early life “injury” could permanently alter nociceptor gene expression and subsequent activity, essentially priming the nervous system to subsequent damage^59^. In practice, this concept known as hyperalgesic priming^60,61^, manifests as exacerbated responses to normally sub-threshold stimuli, but only in tissues or organisms that were previously injured. In viewing SCD through this lens, chronic daily SCD pain may be largely attributed to this phenomenon. Exposure to noxious hemolysis products during critical early-life periods may prime the peripheral nervous system such that exposure to normally innocuous stimuli (*i.e*., mechanical force associated with normal blood flow, minimally elevated chemokine levels in circulation) induce widespread pain later in life. If true, this finding would provide even more support for early and aggressive disease management in individuals born with SCD.

In conclusion, these studies are the first to demonstrate a pediatric pain phenotype in SCD mice. Additionally, this work systematically determined that life-long HU use decreases chronic SCD pain by reducing innate immune system tone. Coupled with the pilot QST and pain diary studies completed in patients^12,14^, these data provide compelling evidence that chronic HU use may not only alleviate acute pain in individuals with SCD but also prevent the development of severe chronic SCD pain.

## Acknowledgements

This work was supported by grants from the National Institutes of Health (grant R00HL155791 to K.E.S.) and the Rita Allen Foundation (Award in Pain to K.E.S.). The authors acknowledge Dr. Lena Nguyen for assistance with western blot analysis, Dr. Benedict Kolber and lab for access to ChemiDoc imaging systems, and Dr. Ted Price and lab for plate reader access.

## Authorship

Designed research: NJG, KES Performed research: NJG, SRR

Contributed vital new reagents or analytical tools: LERF, MDB, HM

Collected data: NJG, SRR, ZM, KES

Analyzed and interpreted data: NJG, SRR, ANP, KES Performed statistical analysis: NJG, KES

Writing manuscript: NJG, KES Editing manuscript: NJG, AMB, KES Funding: KES

Conflict of interest disclosure: The authors declare no competing financial interests.

